# Tumor-induced alterations in single-nucleus transcriptome of atrophying muscles indicate enhanced protein degradation and reduced oxidative metabolism

**DOI:** 10.1101/2023.10.26.564119

**Authors:** Samet Agca, Aylin Domaniku Waraich, Sevval Nur Bilgic, Melis Sucuoglu, Meric Dag, Sukru Anil Dogan, Serkan Kir

## Abstract

Tumor-induced skeletal muscle wasting in the context of cancer cachexia is a condition with profound implications for patient survival. The loss of muscle mass is a significant clinical obstacle and is linked to reduced tolerance to chemotherapy and increased frailty. We investigated muscle gene expression at single nucleus level in cachectic mice and revealed distinct myonuclear gene signatures and a shift towards type IIb myonuclei. Notably, atrophy-related genes, including *Atrogin1*, *MuRF1* and *Eda2r* were upregulated in these myonuclei, emphasizing their crucial role in muscle wasting. Activation of the Ectodysplasin A2 Receptor (EDA2R) pathway suppressed gene sets related to muscle contraction and oxidative metabolism, indicating its involvement in transcriptional reprogramming. Our study also highlighted the negative impact of tumors on oxidative metabolism in muscle tissue and their influence on the transcriptomes of mononuclear cells in skeletal muscle. These findings contribute to a deeper understanding of the molecular mechanisms underlying cancer cachexia.

## Introduction

Tumor-induced skeletal muscle wasting is one of the characteristics of cancer cachexia, a deadly condition that negatively influences prognosis and the survival of patients. Progressive weight loss due to the wasting of muscle and adipose tissues affects the majority of patients with gastric, pancreatic, lung, colorectal and prostate cancers and accounts for at least 20% of all cancer deaths.^1^ Muscle wasting is associated with frailty, chemotherapy intolerance and lack of response to treatment.^2^ Wasting is driven by the atrophy of muscle tissue, which involves reduced protein synthesis and excessive proteolysis.^3^ Loss of muscle mass results in impaired physical strength and the consequent poor quality of life that is often irreversible due to lack of effective therapeutics.^4^ Research oriented toward understanding molecular mechanisms underlying muscle atrophy is essential for developing new tools for treatment.

Our recent work identified the Ectodysplasin A2 Receptor (EDA2R) as a potential marker and therapeutic target for muscle atrophy.^5^ Gene expression analysis in muscle samples from tumor-bearing mice and cachectic cancer patients demonstrated upregulated *EDA2R* levels. Activation of EDA2R by its ligand, A2 isoform of EDA or EDA-A2, caused atrophy in cultured myotubes and the muscle tissue of mice, which involved elevated expression of muscle atrophy markers *Atrogin1* and *MuRF1*.^5^ The depletion of EDA2R or its downstream signaling prevented the loss of muscle mass and function in tumor-bearing mice, underscoring the prominent role of the EDA2R pathway in muscle wasting.^5^ We originally identified *Eda2r* upregulation in the atrophying muscles using whole transcriptome analysis, which has been instrumental in studying disease states in different tissues. However, a major limitation of this technique is that expression profiles of a heterogeneous cell population is averaged and valuable information on differential gene expression in specific subsets of the cells is lost. In this study, we utilized high-throughput, high-resolution transcriptomics to determine tumor-induced alterations in muscle gene expression at single nucleus resolution.

Skeletal muscle tissue is composed of giant syncytial cells called myofibers, each containing numerous myonuclei. While single-cell RNA sequencing (scRNA-seq) enabled studying mononucleated cells in muscle tissue,^6^ high-throughput profiling of both mononuclear and multinuclear cells is achieved by single-nucleus (sn)RNA-seq. This technique involves nuclear isolation and detects nuclear-enriched transcripts, including pre-mRNA.^7,8^ snRNA-seq studies on skeletal muscle tissues indicated that *Titin*-expressing myonuclei account for the majority of the nuclei (∼65%) while the remaining originates from mononuclear cells, including muscle satellite cells (MuSC), fibro-adipogenic progenitors (FAPs), endothelial cells, tenocytes, smooth muscle cells and immune cells.^7–16^ Adult skeletal muscle comprises fast and slow twitch myofibers classified by the expression of specific myosin heavy chain (MYH) genes. In mice, slow myofibers express *Myh7* (type I) while fast myofiber types express *Myh2* (type IIa), *Myh1* (type IIx) or *Myh4* (type IIb). snRNA-seq analysis enables the detection of distinct myonuclear gene signatures representing myofiber types and specialized functions, such as the neuromuscular junction (NMJ) and the myotendinous junction (MTJ).^17,18^ Here, we report that remote tumor growth in mice impacted the transcriptome of muscle tissue, which exhibited the enrichment of type IIb myonuclear gene signatures, the activation of muscle atrophy-related mechanisms, including EDA2R, and the suppression of pathways associated with muscle contractility and oxidative metabolism.

## Results

### snRNA-seq analysis of atrophying muscles identifies mononuclear and myonuclear gene signatures

We utilized Lewis Lung Carcinoma (LLC) cells to induce cachexia and muscle atrophy in mice. Syngeneic C57BL/6 mice injected with LLC cells were sacrificed 16 days after tumor inoculation while they experienced moderate cachexia and loss of muscle mass and function.^5^ We investigated single-nucleus transcriptomes of the tibialis anterior (TA) muscle from tumor-bearing mice and their non-tumor-bearing controls. TA muscles of the cachectic mice exhibited evident wasting as the tissue weight and average muscle fiber cross-sectional area were reduced significantly in this group (Figure 1A-C). Muscle fibers with a small cross-sectional area were enriched (Figure 1D). Muscle samples from 6 different mice in each group were pooled for nuclear isolation. DAPI-stained nuclei were subjected to fluorescence-activated sorting. We constructed single-nucleus libraries using 10X Genomics applications following the manufacturer’s guidelines and performed RNA sequencing. We utilized the Cell Ranger software and Seurat R package to perform quality control and data analysis as detailed in the methods section.

**Figure 1.**
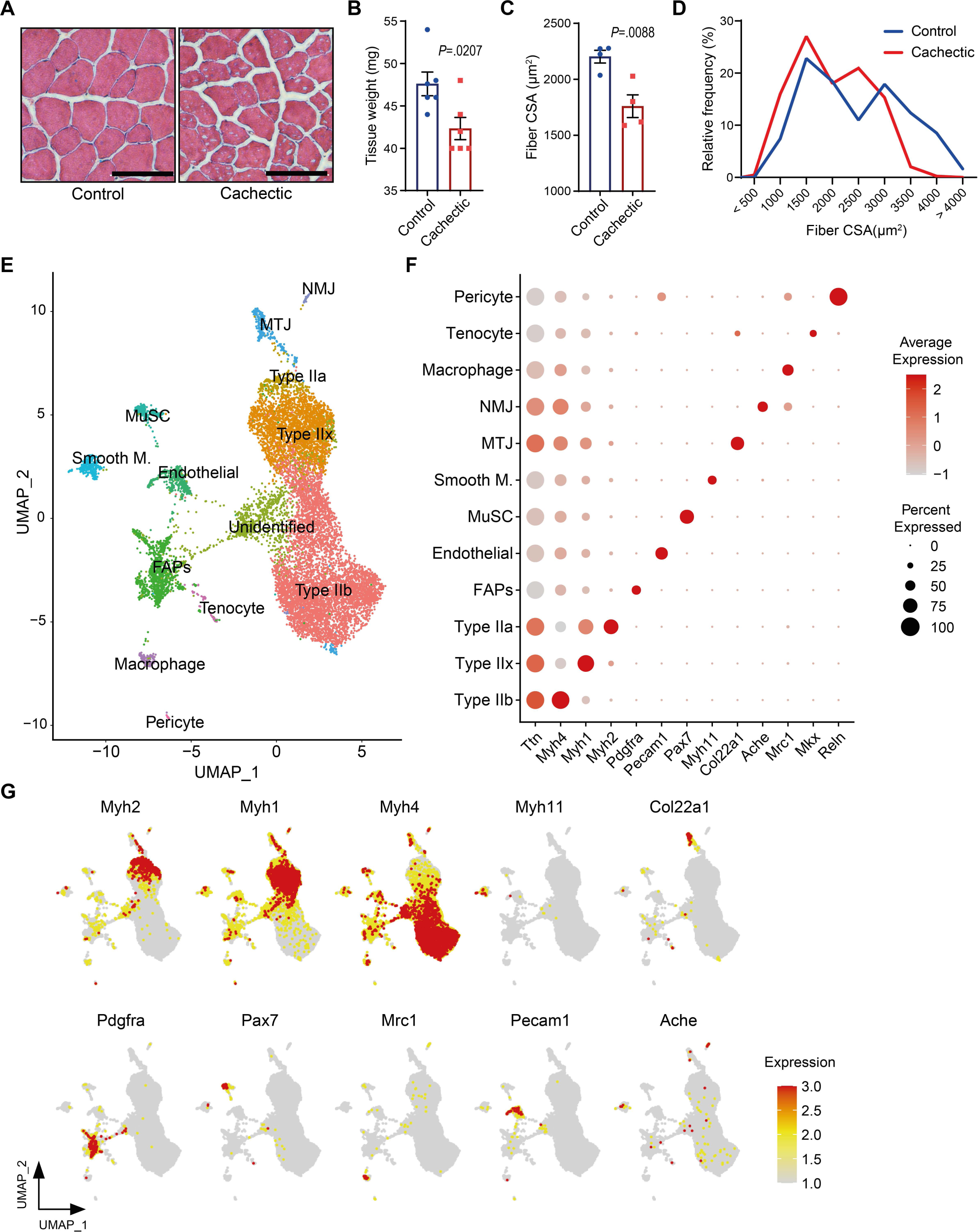
snRNA-seq analysis of atrophying muscles identifies mononuclear and myonuclear gene signatures. (A) Hematoxylin and eosin staining of the tibialis anterior (TA) muscle cross-sections. Scale bars, 100 µm. (B) TA muscle weight of control and cachectic mice. (n=6) (C) Comparison of mean cross-sectional areas of TA muscle fibers (n=4) and (D) distribution of fiber frequencies. (E) Uniform manifold approximation and projection (UMAP) visualization of nuclear clusters; each distinct cell type is color-coded. (F) Dot plot of marker gene expression for each nuclear cluster. The size of the dots represents the percentage of nuclei expressing the marker gene and the red color intensity indicates the expression level. (G) UMAP plots of marker gene expression of each nuclear cluster. (B,C) Unpaired two-sided Student’s t-test was used for statistical analysis. Data are represented as individual points and mean ± SEM.

We analyzed a total number of 12.335 nuclei, comprising 6.422 nuclei from the control group and 5.892 nuclei from atrophying muscles. Using the Seurat and Uniform Manifold Approximation and Projection (UMAP) clustering, we identified 22 distinct unsupervised clusters (Figure S1A). Based on the expression of specific signature genes (Figure S1B and S1C), all clusters were assigned to 12 known cell types, consisting of type II myofibers (including IIa, IIb and IIx), MTJ, NMJ, FAPs, endothelial cells, MuSC, smooth muscle cells, macrophages, tenocytes and pericytes, and unidentified nuclei (cluster 4) (Figure 1E and 1F). A heatmap of the top 5 markers of the identified clusters demonstrated distinct gene expression patterns between these groups (Figure S1C). UMAP clustering of signature genes, including *Myh2* (type IIa), *Myh1* (type IIx), *Myh4* (type IIb), *Myh11* (smooth muscle), *Col22a1* (MTJ), *Pdgfra* (FAPs), *Pax7* (MuSC), *Mrc1* (macrophages), *Pecam1* (endothelial cells), *Ache* (NMJ), illustrated the distribution of these clusters (Figure 1G).

### Tumors induce the enrichment of type IIb myonuclear signatures in TA muscle

A comparison of nuclear populations between the control and cachectic muscles indicated that the distribution of these clusters was altered in the presence of tumors (Figure 2A). The predominant cell type in the control TA muscles is type II fibers containing 64.86% of all nuclei. The representation of type II myonuclei increased to 76.22% in the cachectic muscles. An increase in the proportion of type IIb myonuclei from 41.03% to 52.75% accounted for the bulk of this change (Figure 2B). In addition, slightly increased proportions of NMJ and MTJ myonuclei were detected. Interestingly, the representation of mononuclear cells, including FAPs, endothelial cells, smooth muscle cells, macrophages, tenocytes and pericytes, was reduced in the cachectic muscles (Figure 2B). These findings argue that tumor growth induced the enrichment of myonuclear signatures and particularly type IIb myonuclei in muscle tissue.

**Figure 2.**
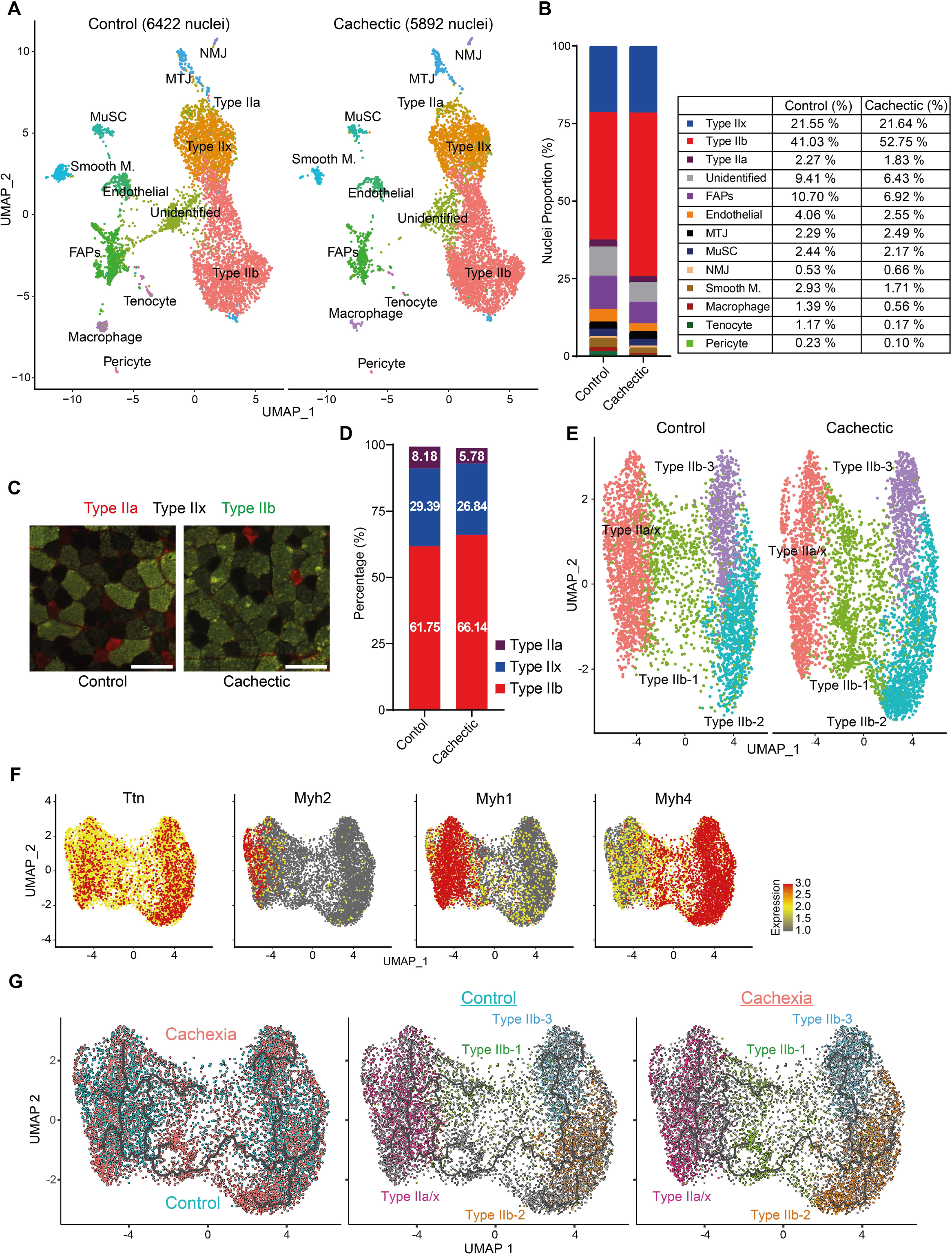
Tumors induce the enrichment of type IIb myonuclear signatures in TA muscle. (A) UMAP plot of nuclear clusters in control (left panel) and cachectic (right panel) TA muscles, each distinct cell type is color-coded. (B) Proportions of nuclear clusters in control and cachectic muscles, relative to the total number of nuclei in each group. (C) Representative images of immunofluorescence (IF) staining of TA muscle sections. Green and red signals visualize Myh4 (type IIb) and Myh2 (type IIa) expression, respectively, while dark fibers are type IIx. Scale bars, 100 µm. (D) Percentages of specific fiber types in IF staining of TA muscle sections. (E) UMAP plot of myonuclear clusters in control (left panel) and cachectic (right panel) TA muscles. (F) UMAP plots representing levels of marker gene expression in myonuclei. (G) UMAP plots depicting the trajectory analysis of myonuclei from control and cachectic muscles. The myonuclei are colored according to conditions in the left panel and according to identified subtypes in the middle and the right panel. Data are represented as individual points.

We analyzed the change in the composition of myofibers to investigate the impact of the enrichment of type IIb myonuclear signatures upon tumor inoculation. Immunofluorescence staining of TA muscle sections showed that the majority of muscle fibers are type IIb and type IIx while type IIa and type I fibers make up a small fraction (Figure 2C). The percentage of type IIb fibers increased from 61.59 to 66.11 in the cachectic muscles arguing that the tumor-induced increase in the proportion of type IIb myonuclei also reflected in the frequency of these myofibers (Figure 2D). We further analyzed type II myonuclear gene signatures in our snRNA-seq dataset and identified 4 distinct myonuclear populations (Figure 2E). We categorized them based on *Myh* gene expression and detected a cluster representing type IIa and type IIx myonuclei (type IIa-x) and 3 distinct clusters representing type IIb myonuclei (type IIb-1, IIb-2 and IIb-3) (Figure 2F and S2A). Upon cachexia, a 10% drop in the number of type IIa-x myonuclei and about 5% increases in each of type IIb-1 and type IIb-2 myonuclei were detected (Figure S2B). Trajectory analysis of type II myonuclei revealed a continuous path originating from type IIa-x myonuclei, extending into type IIb-1 and branching out to type IIb-2 and type IIb-3 clusters, implying that cancer cachexia promoted the transition of type IIa and type IIx myonuclei towards the type IIb identity (Figure 2G). However, it cannot be ruled out that remote tumor growth also induces the fusion of myogenic progenitors which contribute myonuclei with type IIb myonuclear characteristics.

### Tumors promote the enrichment of atrophy-related myonuclear signatures in TA muscle

We next investigated differentially expressed genes (DEGs) in myonuclei. Comparison of control and cachectic muscles indicated upregulated expression of atrophy-related genes in the latter group, including *Atrogin1* (*Fbxo32*) and *MuRF1* (*Trim63*) (Figure 3A and 3B). These two genes encode E3 ubiquitin ligases that are well-known inducers of muscle protein breakdown and atrophy.^3^ The expression of *Atrogin1* and *MuRF1* was elevated in all myofiber types but this change was particularly striking in type IIb myonuclei (Figure 3B and S3A), arguing a predominant effect on type IIb fibers during muscle wasting. Other E3 ubiquitin ligase-associated genes linked to protein degradation, such as *Asb2* and *Klhl38* were also upregulated (Figure 3B).^19,20^ Notably, expression of genes involved in the regulation of metabolic activities, including *Pik3r1*, *Pdk4*, *Foxo1*, *Rorc* and *Lpin1*, increased in type IIb myonuclei upon cachexia (Figure 3B). Particularly, *Pdk4*, which encodes a kinase that phosphorylates pyruvate dehydrogenase and inhibits pyruvate oxidation, and the transcription factor *Foxo1* stood out as muscle atrophy-related markers.^21,22^ Genes downregulated in the cachectic type IIb myonuclei include *Myh1*, *Myl1*, *Acta1*, *Actg1*, *Ank2*, *Fhl1*, *Mylpf*, *Fhod3* and *Mybpc2*, which are involved in muscle contraction and its regulation (Figure 3B). Notably, the expression of creatine kinase genes *Ckm* and *Ckmt2*, and the muscle-specific glycolytic enzyme *Eno3* was also reduced, implicating that energy metabolism supporting muscle contraction was negatively impacted by tumors (Figure 3B). Genes involved in positive regulation of muscle mass, including *Fhl1* and polyamine biosynthesis enzymes *Amd1* and *Smox* were also downregulated (Figure 3B).^23–25^ Up- or down-regulation of the majority of these genes was more pronounced in type IIb myonuclei compared to type IIx and type IIa myonuclei. Type IIb-enriched gene expression changes were validated by RT-qPCR analysis of bulk RNA from TA muscle samples (Figure 3D). Distinct gene expression patterns were also detected in type IIx and type IIa myonuclei (Figure S3B). For instance, transcription factors *Foxp2* and *Ppara* were particularly up- and down-regulated, respectively, in type IIx myonuclei while muscle proteins nebulin (*Neb*) and tropomyosin-1 (*Tpm1*) were suppressed in type IIa myonuclei (Figure S3B). Upon tumor growth, *Atrogin1* expression was enriched in specialized MTJ and NMJ myonuclei, which also exhibited distinctive changes in their transcriptomes, including increased *Eda* expression in MTJ and reduced *Deptor* and *Nos1* expression in NMJ (Figure S3C and S3D). Additionally, we identified increased expression of *Eda2r*, which showed marked upregulation in the cachectic type IIb myonuclei, a pattern similar to *Atrogin1* and *MuRF1* (Figure 3B, 3E and S3A). Mild enrichment in *Eda* mRNA is also detectable in type II myonuclei (Figure S3A). Our recent work demonstrated that the EDA2R signaling mediates tumor-induced *Atrogin1* and *MuRF1* expression in muscle tissue, placing EDA2R at upstream of these atrophy markers.^5^

**Figure 3.**
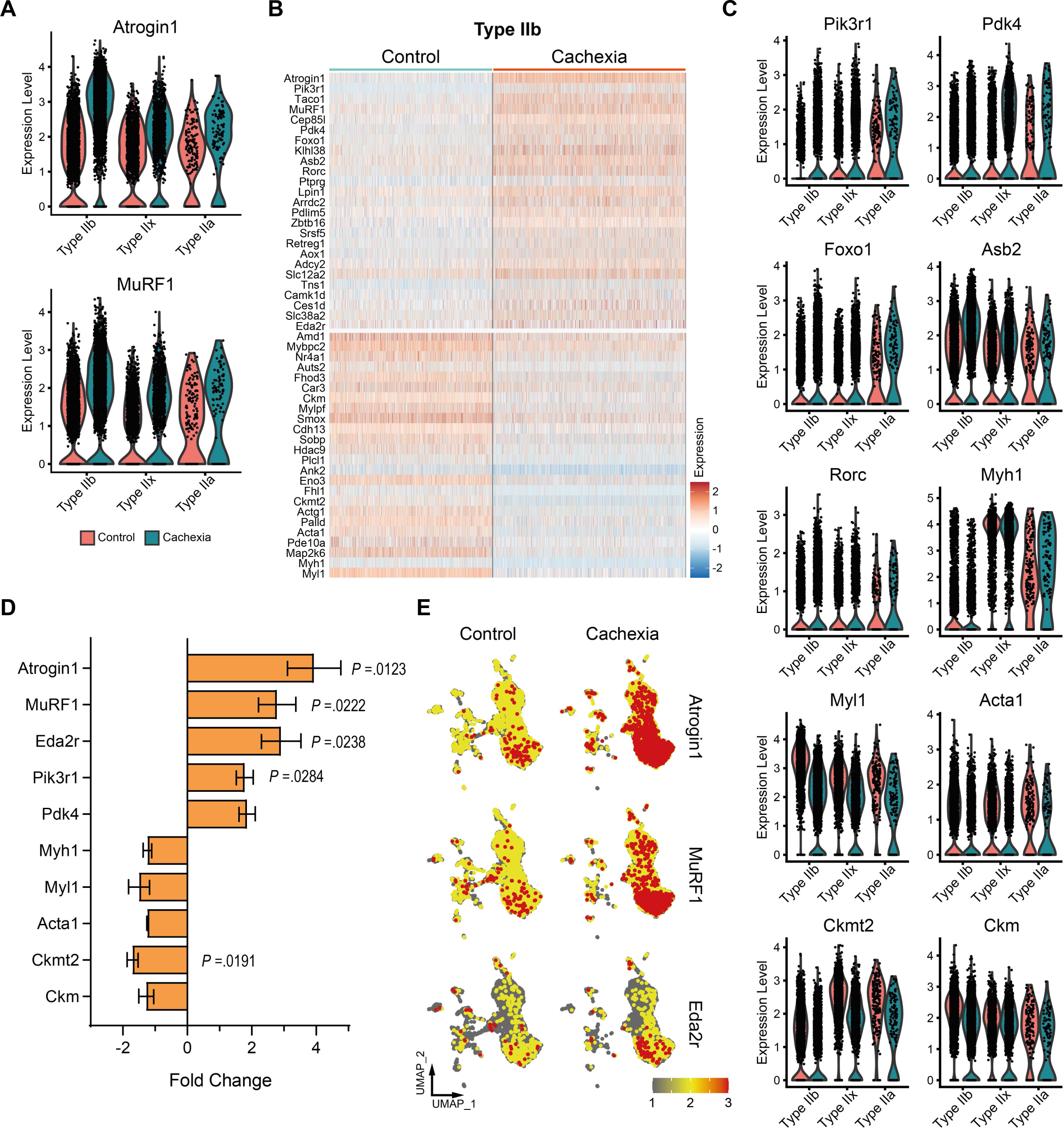
Tumors promote the enrichment of atrophy-related myonuclear signatures in TA muscle. (A) Violin plots of *Atrogin1* and *MuRF1* expression levels in myonuclei (red: control, blue: cachectic). (B) Heatmap of the differentially expressed genes (DEGs) in type IIb myonuclei. (C) Violin plot of the expression levels of selected genes in myonuclei (red: control, blue: cachectic). (D) RT-qPCR analysis of selected genes in bulk RNA isolated from control and cachectic TA muscle samples. (n=4) (E) UMAP plots representing gene expression levels in control and cachectic nuclei. (D) Unpaired two-sided Student’s t-test was used for statistical analysis. Data are represented as individual points or mean ± SEM.

### EDA2R activation and tumor inoculation suppress gene sets related to muscle contraction and oxidative metabolism

Previously, we described muscle atrophy-inducing effects of the EDA2R pathway.^5^ Stimulation of mouse primary myotubes with EDA-A2 promoted cellular atrophy detected by myosin heavy chain immunostaining (Figure 4A). EDA-A2 administration significantly reduced myotube diameter (Figure 4B). To investigate global transcriptional changes driven by EDA2R activation, we performed bulk RNA sequencing. Gene set enrichment analysis (GSEA) of DEGs revealed that hallmark gene sets involving interferon response, inflammatory response, TNFα signaling via NFĸB, IL6-JAK-STAT signaling and TGFβ signaling were enriched in EDA-A2-treated myotubes (Figure 4C and S4A). The roles of these pathways and inflammation in cancer cachexia has been well-established.^26^ Gene sets involving myogenesis, oxidative phosphorylation, fatty acid oxidation and angiogenesis were enriched in the control group compared to the EDA-A2-treated samples (Figure 4C). Because *Eda2r* expression was induced in type IIb myonuclei of cachectic muscles, we performed GSEA of these myonuclei using hallmark gene sets and compared it with the analysis of the EDA-A2 dataset. Remarkably, pathways activated by EDA-A2, such as NFĸB, JAK-STAT and TGFβ, were also enriched in the cachectic type IIb myonuclei while pathways suppressed by EDA-A2, including myogenesis, oxidative phosphorylation and fatty acid oxidation, behaved similarly in the cachectic type IIb myonuclei (Figure 4D, S4B and S4C). The overlap between the gene set enrichment of EDA-A2-treated myotubes and type IIb myonuclei argues that EDA2R upregulation in the cachectic myonuclei likely contributes to the transcriptional reprogramming taking place after tumor inoculation.

**Figure 4.**
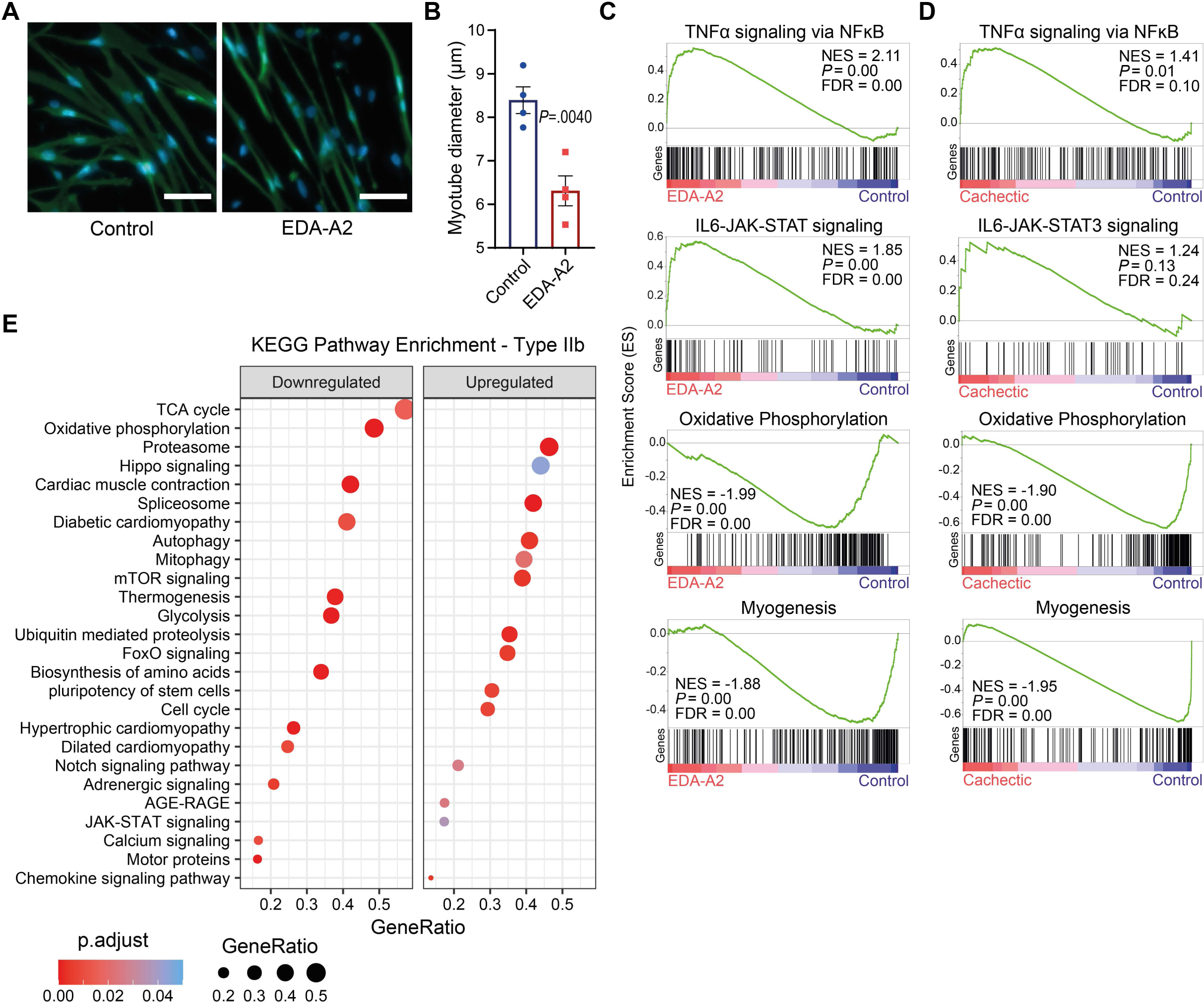
EDA2R activation and tumor inoculation suppress gene sets related to muscle contraction and oxidative metabolism. (A) Myosin heavy chain immunofluorescence staining of control (left panel) and recombinant EDA-A2-treated (right panel) mouse primary myotubes. Scale bars, 50 µm. (B) Comparison of average myotube diameters between conditions (n=4). (C,D) Gene set enrichment (GSEA) plots of hallmark gene sets enriched in EDA-A2-treated myotubes (C) and cachectic type IIb myonuclei (D). NES, normalized enrichment score; FDR, false discovery rate. (E) KEGG pathways enriched in cachectic type IIb myonuclei. The size of the dots represents the enriched gene ratio in each pathway and the red color intensity represents adjusted p values. (B) Unpaired two-sided Student’s t-test was used for statistical analysis. Data are represented as individual points and mean ± SEM.

We next performed further GSEA analysis of our data using KEGG gene sets. In type IIb myonuclei, gene sets including proteasome, ubiquitin-mediated proteolysis, autophagy, FOXO signaling, stem cell pluripotency, and hippo signaling were upregulated in response to cachexia while gene sets such as motor proteins, cardiac muscle contraction, cardiomyopathies, glycolysis, oxidative phosphorylation, TCA cycle and thermogenesis were downregulated (Figure 4E). Remarkably, most gene sets suppressed in cachectic myonuclei were also downregulated in myotubes treated with EDA-A2, arguing that the EDA2R pathway activation may be related to reduced expression of genes involved in muscle contractility and metabolism (Figure S4C). Gene set enrichment patterns similar to that of type IIb myonuclei were also detected in type IIx, type IIa, MTJ and NMJ myonuclei (Figure S4D-S4H), demonstrating that muscle wasting signatures are not restricted to type IIb myonuclei. However, enrichment of distinct gene sets was also identified in non-type IIb myonuclei, including Wnt, PPAR and AMPK signaling pathways. Notably, the gene set for spliceosome was enriched in all myonuclei upon cachexia, implicating that this process may have an underappreciated role in muscle loss (Figure 4E and S4D-S4F). Serine and arginine rich splicing factor 5 (*Srsf5*) is particularly upregulated in myonuclei (Figure 3B).

### Muscle oxidative metabolism is suppressed by tumors and EDA2R activation

Intrigued by the suppression of the gene sets for oxidative phosphorylation and thermogenesis in myotubes treated with EDA-A2 and myonuclei of tumor-bearing mice, we investigated O2 consumption of the muscle cells. Upon measuring basal and maximal O2 consumption rates in primary myotubes, we detected a clear trend for reduced oxidative metabolism after the administration of recombinant EDA-A2 protein (Figure 5A-B). We then isolated mitochondria from the TA muscles of control and tumor-bearing mice and measured their O2 consumption rates. Oxygraphic analysis of isolated mitochondria supplemented with both Complex I- and Complex II-linked substrates under phosphorylating, non-phosphorylating, and uncoupled conditions demonstrated decreased mitochondrial respiration in the atrophying muscles (Figure 5C). Next, we quantified reactive oxygen species (ROS) generation in skeletal muscle mitochondria by measuring hydrogen peroxide (H2O2) production. Mitochondrial ROS production was similar in these samples (Figure 5D). In agreement with these findings, metabolic pathway enrichment analysis indicated that processes including oxidative phosphorylation, TCA cycle and glycolysis were downregulated in all type II myonuclei whereas pyruvate metabolism and fatty acid degradation were suppressed in type IIb and type IIx myonuclei (Figure 5E). Pathways for the metabolism of amino acids, such as alanine, aspartate, glutamate, arginine, proline, valine, isoleucine, isoleucine, tryptophan, phenylalanine, tyrosine, cysteine and methionine were particularly suppressed in type IIb myonuclei (Figure 5E). In contrast, pathways including N-glycan biosynthesis and glutathione metabolism were enriched in type II myonuclei (Figure 5E). Since type IIb and type IIx myonuclei are more numerous in TA muscles, their metabolic activities are expected to be rather predominant. Overall, our findings indicate that tumor-induced muscle wasting is associated with reduced oxidative metabolism, including the metabolism of most amino acids in this tissue.

**Figure 5.**
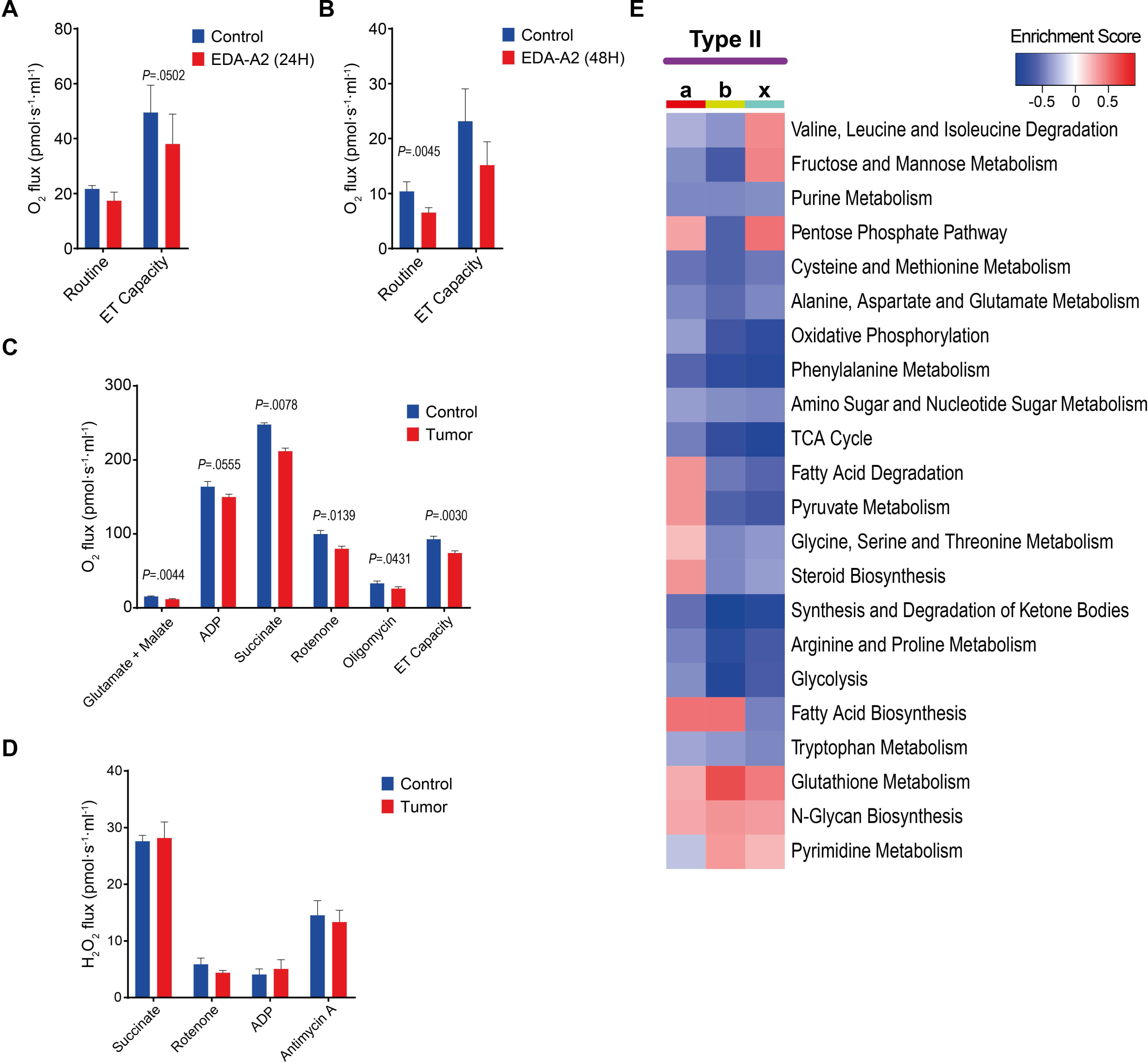
Muscle oxidative metabolism is suppressed by tumors and EDA2R activation. (A,B) O2 consumption rates in primary myotubes treated with EDA-A2 for 24 hours (A) and for 48 hours (B). Routine represents the basal respiratory activity, and “Electron Transfer (ET) capacity” is the uncoupled respiration obtained by CCCP titration (n=3). (C,D) O2 consumption (C) and H2O2 production (D) rates in mitochondria isolated from TA muscles of control and tumor-bearing mice (n=4). (E) Heatmap of metabolic pathway enrichment analysis in type II myonuclei. Paired two-sided Student’s t-test was used for statistical analysis. Data are represented as individual points and mean ± SEM.

### Tumors alter the transcriptomes of mononuclear cells in muscle tissue

We also investigated differential gene expression in the nuclei of mononuclear cells of skeletal muscle. The most numerous mononuclear cells; FAPs, endothelial and smooth muscle cells exhibited striking expression patterns. Although tumor-induced suppression of gene sets associated with oxidative phosphorylation and cardiomyopathy, and the enrichment of gene sets associated with the proteasome, spliceosome, JAK-STAT signaling and FOXO signaling are reminiscent of myonuclear transcriptional changes, FAPs gene signatures also involved the upregulation of gene sets for amino acid degradation, ferroptosis and adipocytokine signaling, and downregulation of gene sets for focal adhesion and extracellular matrix (ECM)-receptor interaction (Figure S5A). Notably, the heatmap of transcripts with the highest expression changes in FAPs was populated by genes with extracellular matrix-related functions, including *Nid1*, *Fap*, *Lum*, *Sparc*, *Cd34*, *Fbn1*, *Itgbl1*, *Mxra7*, which were suppressed in the cachectic muscles (Figure S5B). Gene expression analysis in endothelial and smooth muscle cells of muscle vasculature also revealed the suppression of gene sets associated with oxidative metabolism in response to tumors (Figure S5C and S5D). Therefore, it is likely that reduced O2 consumption is relevant for both myofibers and the majority of mononuclear cells in the cachectic muscle. Upon tumor growth, genes involved in angiogenesis and vascular growth, such as *Vegfa* and *Fhl1*, were suppressed in smooth muscle cells while *Hspg2* (perlecan) and *Ets1* were downregulated in endothelial cells (Figure S5E and S5F).^27–29^ In addition, muscle atrophy-related genes, *Osmr* and *Stat3* were upregulated in endothelial cells (Figure S5F). Notably, KEGG gene sets for lipid and pro-atherosclerosis, and adipocytokine signaling were enriched in endothelial cells of the cachectic muscles whereas fluid shear stress and atherosclerosis (mostly anti-atherosclerotic factors), and focal adhesion gene sets were suppressed (Figure S5C). Gene signatures in smooth muscle cells in the cachectic muscles exhibited the enrichment of gene sets, including Notch signaling, adherens junction, focal adhesion, vascular smooth muscle contraction, cGMP-PKG signaling, and autophagy, implying that these cells undergo significant remodeling under cachectic conditions (Figure S5D).

## Discussion

The syncytial nature of myofibers is a major obstacle against a high-resolution analysis of gene expression in skeletal muscle. scRNA-seq is not suitable for studying myonuclear transcriptomes since this technique detects very few muscle cells.^30^ However, snRNA-seq is a reliable alternative, which identifies both myonuclear and mononuclear transcriptomes in muscle tissue.^7,8^ It should be noted that snRNA-seq only detects nuclear-enriched transcripts, including pre-mRNA, and therefore alterations in the cytoplasmic mRNA pool remain hidden. Since cytoplasmic transcripts are unaccounted for, the resolution is not optimal for genes with low expression levels. Despite these limitations, snRNA-seq analysis of muscle tissue has been used to document the impact of disease states, such as Duchenne muscular dystrophy and denervation, on the transcript levels.^10,12,14^ In this study, we utilized snRNA-seq to study the cachectic skeletal muscle of tumor-bearing mice. Our results revealed that remote tumor growth transforms gene expression in mononuclear cells and myonuclei. Analysis of differentially expressed genes indicated the adoption of myonuclear gene signatures associated with muscle atrophy-related processes, such as enhanced protein degradation and reduced oxidative metabolism.

snRNA-seq analysis enables the determination of myofiber type-specific gene signatures in myonuclei. We demonstrated that type IIb myonuclei were enriched in the cachectic muscles. From these data, it is unclear if cachexia induced the emergence of new type IIb myonuclei or the adoption of a type IIb-resembling signature in the existing myonuclei via transcriptional reprogramming. However, pseudotime trajectory analysis indicated that a transition from type IIa-x myonuclei towards the type IIb identity is possible. Interestingly, the increase in the proportion of type IIb myonuclei is also reflected in the frequency of type IIb myofibers. In agreement with previous studies which reported a transition towards type IIb myofibers in cachectic muscles,^31–33^ a slight increase in the percentage of these myofibers was also detected in the tibialis anterior (TA). Whether tumor-induced enrichment of type IIb myonuclear signatures is limited to TA muscle remains to be determined. Additional studies should address how myonuclear identity and gene expression profiles change in different muscle tissues in response to cancer cachexia.

Profound upregulation of atrophy-related genes, including E3 ubiquitin ligases *Atrogin1* and *MuRF1*, transcription factor *Foxo1*, and pyruvate dehydrogenase kinase *Pdk4* was detected in myonuclei isolated from the cachectic muscles.^21,22^ While *Asb2* as a negative regulator of muscle mass was induced in these samples, positive regulators of muscle mass, such as *Fhl1*, *Amd1* and *Smox* were downregulated.^19,23–25^ This was accompanied by reduced levels of genes involved in muscle contraction and energy metabolism, including *Myh1*, *Myl1*, *Acta1*, *Eno3*, *Ckm* and *Ckmt2*. Notably, differential gene expression related to muscle atrophy and metabolism was more pronounced in type IIb myonuclei, in which *Eda2r* is also upregulated. EDA2R signaling has recently been described to play a prominent role in tumor-induced muscle wasting.^5^ It is likely that EDA2R upregulation in the cachectic myonuclei contributes to the transcriptional reprogramming taking place after tumor inoculation. In fact, gene set enrichment analysis demonstrated that EDA2R activation and tumor inoculation led to similar expression patterns in muscle cells including the stimulation of NFĸB, JAK-STAT and TGFβ pathways and the suppression of myogenesis, oxidative phosphorylation and fatty acid oxidation. Our flux measurements indicated reduced O2 consumption in myotubes upon the activation of EDA2R signaling. In agreement with a previous study,^34^ mitochondrial respiration was also suppressed in cachectic muscles. Reduced mitochondrial oxidation and inefficient metabolism potentially support muscle wasting. However, elevated energy expenditure and O2 consumption were previously described in tumor-bearing mice, which were attributed to enhanced adipose tissue browning.^35,36^ Our findings argue that skeletal muscle metabolism with reduced oxidative potential is unlikely to contribute directly to the accelerated metabolic rate prevalent in the cachectic mice.

snRNA-seq analysis of mononuclear cells in the cachectic muscles also revealed substantial transcriptional changes, including the downregulation of genes with extracellular matrix-related functions in FAPs and the downregulation of genes involved in vascular growth in smooth muscle and endothelial cells. The suppression of gene sets associated with oxidative metabolism in these cells argues that impairment in mitochondrial respiration is not restricted to myofibers and this effect may stem from a deficiency in the vasculature of the cachectic muscle tissue. In fact, a recent study utilizing cachexia-inducing tumor models described weakened vascular functions in atrophying muscles.^37^ In skeletal muscle tissue, tumors likely exert an extensive impact that does not merely target myofibers. Finally, gene sets identified in this study contribute significantly to our understanding of potential therapeutic targets for mitigating cancer cachexia and associated muscle wasting. Future research should focus on uncovering the mechanisms involving these genes and their specific roles in muscle wasting.

## Acknowledgments

We gratefully acknowledge the use of the Koc University Research Center for Translational Medicine (KUTTAM) animal facility infrastructure. This work was supported by the EMBO installation grant (#4162) to S. Kir

## Author contributions

S.K. conceived and designed the experiments. S.A., A.D., S.N.B., M.S., M.D., S.A.D, and S.K. performed the experiments. S.A., A.D., S.N.B., M.S. and S.A.D. analyzed the data. S.A., S.A.D and S.K. wrote the manuscript.

## Declaration of interests

The authors declare no competing interests.

## Methods

### Mice

Mice were housed at 22 °C and under 50% humidity with 12 hours of light and 12 hours of dark cycles (07:00 – 19:00) and given ad libitum access to a standard rodent chow diet and water in Koc University Animal Research Facility in accordance with institutional policies and animal care ethics guidelines. 8-12 weeks old male mice with C57BL/6 background were used for the experiments. Lewis lung carcinoma (LLC) cells were used for tumor inoculation. LLC cells were cultured in DMEM (Sigma, no. 5796) with 10% fetal bovine serum (FBS) and penicillin/streptomycin (Invitrogen). 5 × 10^6^ LLC cells were injected subcutaneously over the flank while control mice received PBS only. Muscle tissues were harvested at 16 days after LLC inoculation.

### Tissue histology

Isopentane was cooled with liquid nitrogen until it was at least half-frozen. Muscle samples were placed in the cooled isopentane for 10 seconds. The frozen tissues were embedded in Tissue-Tek OCT freezing medium (Sakura) and cut into 8-µm-thick sections using a cryostat. Sections were collected on Superforst Plus slides (Thermo). The sections were fixed with 4% paraformaldehyde and stained with hematoxylin (Merck, no. 105174), 0.1 % hydrochloric acid, eosin (Merck, no. 109844), a 70-100% ethanol gradient, and xylene (Isolab). The cross-sectional area of the muscle fibers was measured using ImageJ software.

### Immunofluorescence staining of muscle sections

Cryosections were incubated with a blocking solution (3% bovine serum albumin + 0.1% Triton X-100 + 5% Horse serum) for 1hr at room temperature and then with monoclonal anti-myosin heavy chain (MyHC) antibodies in the blocking solution overnight at +4°C. MyHC antibodies (DSHB): SC-71 (IgG1, 1:100 dilution) specific for MyHC-2A, BF-F3 (IgM, 1:100 dilution) specific for MyHC-2B, 6H1 (IgM, 1:50 dilution) specific for MyHC-2X and BA-F8 (IgG2b, 1:50 dilution) specific for MyHC-I (slow, alpha and beta fibers). Sections were washed with PBS and incubated in the blocking solution containing goat anti-mouse IgM Alexa Fluor 488 (Abcam, ab150121, 1:500) antibody and goat anti-mouse IgG H&L Alexa Fluor 594 (Abcam, ab150116, 1:500) antibody for 1hr. Finally, the sections were mounted using a homemade mounting medium.

### Primary myoblast culture

Mouse primary myoblasts were isolated from the limb muscles of 2-3 day-old pups and cultured in Ham’s F-10 nutrient mixture (Invitrogen) as described before.^38^ The culture medium was supplemented with 20% fetal bovine serum (FBS, Invitrogen), 2.5 ng/ml of basic fibroblast growth factor (bFGF, Sigma), and penicillin/streptomycin (PS, Invitrogen). For differentiation, cultured myoblast cells were transferred to DMEM (Sigma 5796) with %5 horse serum (Invitrogen) and PS.

### Immunofluorescence staining of primary myotubes

Cells fixed in ice-cold 100% methanol at -20°C for 10 minutes were incubated in a blocking solution containing 3% bovine serum albumin, 0.1% Triton X-100, and 10% horse serum at room temperature for 1 hour. Cells were then incubated with a MyHC antibody (DSHB, MF20) at a 1:1000 dilution in the blocking solution for 1 hour at room temperature. After washing with PBS, the cells were treated with an anti-mouse IgG H&L Alexa Fluor 594 secondary antibody (Abcam, ab150116) at a 1:2000 dilution and DAPI (Cayman) at a 1:3000 dilution in the blocking solution for 1 hour. Cells mounted with a homemade medium were visualized using a Zeiss fluorescence microscope. The diameters of individual myotubes were measured at three different sites with values averaged using Image J software.

### RT-qPCR

Total RNA from cultured myotubes or muscle tissue samples was extracted using Qiazol reagent (Qiagen) and purification was performed using RNA spin columns (Ecotech). TissueLyzer LT (Qiagen) was used to homogenize the tissues. High-capacity cDNA Reverse Transcription kit (Thermo) was used for complementary DNA synthesis. Reactions were set with 25 ng of cDNA, 150 nmol of specific primers and iTaq Universal SYBR Green Supermix (Bio-Rad). Gene expression was analyzed by RT-qPCR using a CFX Connect instrument (Bio-Rad). Calculations for relative mRNA levels were done by ΔΔCt method and normalized to cyclophilin mRNA.

### Bulk RNA-sequencing

Mouse primary myoblasts were differentiated into myotubes for 48 hours and then treated with 250 ng/ml recombinant EDA-A2 for 24 hours. Isolation and DNase treatment of total RNA was performed using Qiazol reagent (Qiagen) and Direct-zol RNA MiniPrep kit (Zymo Research). Isolated samples were prepared with Illumina TruSeq Stranded mRNA Library Prep Kit. Illumina NovaSeq technology was used for pair-end sequencing which produced a total of 60 million (2 × 100bp) paired-end reads per sample. The quality check of raw sequencing data was done with FastQC (v0.11.9) and reports were created with MultiQC (v1.12). Trimming of poor-quality reads and adaptor sequences was performed with trimGalore (v0.6.5). A mouse genome index (GRCm39) was created and reads were mapped using STAR (v2.7.3a). The quality of mapped data was checked with qualimap (v2.2.1). Quantifications of alignments done by HTSeq (v0.11.1).

### Skeletal muscle nuclei isolation

A previously described protocol was used to isolate nuclei.^9^ Briefly, TA muscle samples dissected from 6 mice in each group were combined and processed together. Samples were chopped with dissection scissors and placed into a lysis buffer containing 10 mM Tris-HCl, 10 mM NaCl, 3 mM MgCl2, and 0.1% NP40 in nuclease-free water. Samples were homogenized using a douncer and filtered with 70 µm and 40 µm cell strainers. After centrifugation for 5 min at 500 g at 4 °C, the supernatant was discarded and nuclei were resuspended and stained with DAPI and subjected to fluorescence activated cell sorting.

### Single-nucleus RNA sequencing and bioinformatic analyses

10X Genomics applications were used following the manufacturer’s guidelines (Chromium Next GEM Single Cell 3ʹ Reagent Kits) to prepare the libraries, which were sequenced using the Illumina HiSeq X system. Sequencing data was first analyzed and filtered using Cell Ranger (v7.0) Single-Cell Software Suite provided by 10X Genomics. Data was counted and mapped with cellranger count function with --include-introns option for pre-mRNAs. Further analysis was performed using Seurat (v4.3.0) R (v4.2.2) package on R Studio (v2022.12.0), which filters nuclei, normalizes expression data, and carries out principal component analysis for clustering and Uniform Manifold Approximation and Projection (UMAP) visualization. Seurat R package also performs marker assignment of the clusters and gene ontology analysis of their uniquely expressed genes. ClusterProfiler (4.7.1.003) R package was used for KEGG pathway enrichment analysis and GSEA software (v4.3.2) was used for hallmark gene sets enrichment analysis.

### Mitochondria isolation, oxygen consumption and H2O2 production rate measurements

Mitochondria were isolated from the tibialis anterior muscles. Following dissection, the muscles were cut into small pieces and incubated in ice-cold PBS supplemented with 10 mM EDTA and 0.05% trypsin for 30 min. The supernatant was discarded after centrifugation at 200g for 5 min at 4°C. The pellet was resuspended in IBm1 buffer (67 mM Sucrose, 50 mM KCl, 50 mM Tris-HCl, 10 mM EDTA, 0.2% fatty acid free BSA, pH 7.4) and homogenized using a Teflon-Glass pestle. The homogenous solution was centrifuged at 700g for 10 min at 4°C. The supernatant was transferred into a new tube and centrifuged at 8000g for 10 min at 4 °C. The pellet containing mitochondria was resuspended in 50 μl IBm2 buffer (250 mM Sucrose, 0.3 mM EGTA-Tris, 10 mM Tris-HCl, pH 7.4).

Oxygen consumption rates were measured using an Oroboros O2k-FluoRespirometer (Oroboros Instruments, Innsbruck, Austria). 130 μg of muscle mitochondria was diluted in 2 ml of mitochondrial respiration buffer MiR05 (0.5 mM EGTA, 3 mM MgCl2, 60 mM Lactobionic acid, 20 mM Taurine, 10 mM KH2PO4, 20 mM HEPES, 110 mM Sucrose, 1 mg/ml fatty acid free BSA, pH 7.2) inside the Oroboros O2k-FluoRespirometer chambers. A substrate-uncoupler-inhibitor-titration (SUIT) protocol was used to determine the additive effect of Complex I- and Complex II-linked substrates in isolated mitochondria. The following chemicals are added in order: 10 mM Glutamate, 2 mM Malate, 2.5 mM ADP, 10 mM Succinate, 0.5 µM Rotenone, 10 nM Oligomycin, stepwise 0.5 µM Carbonyl cyanide 3-chlorophenylhydrazone (CCCP), 2.5 µM Antimycin A.

3 × 10^5^ primary myotubes per well were incubated with their culture medium in the Oroboros O2k-FluoRespirometer. Routine respiration represented the physiological coupling state of the myotubes; the “Electron Transfer (ET) capacity” was obtained by stepwise addition of 0.5 µM CCCP.

H2O2 production rate was measured at 37 °C using 130 μg of muscle mitochondria diluted in 2 ml of mitochondrial respiration buffer MiR05 (0.5 mM EGTA, 3 mM MgCl2, 60 mM Lactobionic acid, 20 mM Taurine, 10 mM KH2PO4, 20 mM HEPES, 110 mM Sucrose, 1 mg/ml fatty acid free BSA, pH 7.2) in an Oroboros O2k-FluoRespirometer. The reaction of H2O2 and Amplex® UltraRed (10 µM) is catalyzed by horseradish peroxidase (1 U/ml) and superoxide dismutase (5 U/ml) to produce the red fluorescent compound resorufin. The reaction is initiated by addition of Succinate (10 mM), followed by 0.5 µM Rotenone, 2.5 mM ADP, and 2.5 µM Antimycin A. In between, 0.1 μM H2O2 was added for internal and experimental calibration. The change of emitted fluorescence, which was directly proportional to the H2O2 produced, was calculated.

### Statistical Analysis

Unpaired two-tailed student’s t-test was used for comparisons between the two groups. Paired two-tailed student’s t-test was used for oxygen consumption analysis (Figure 5). Data are represented as mean ± SEM.

### Supplemental information

**Figure S1.**
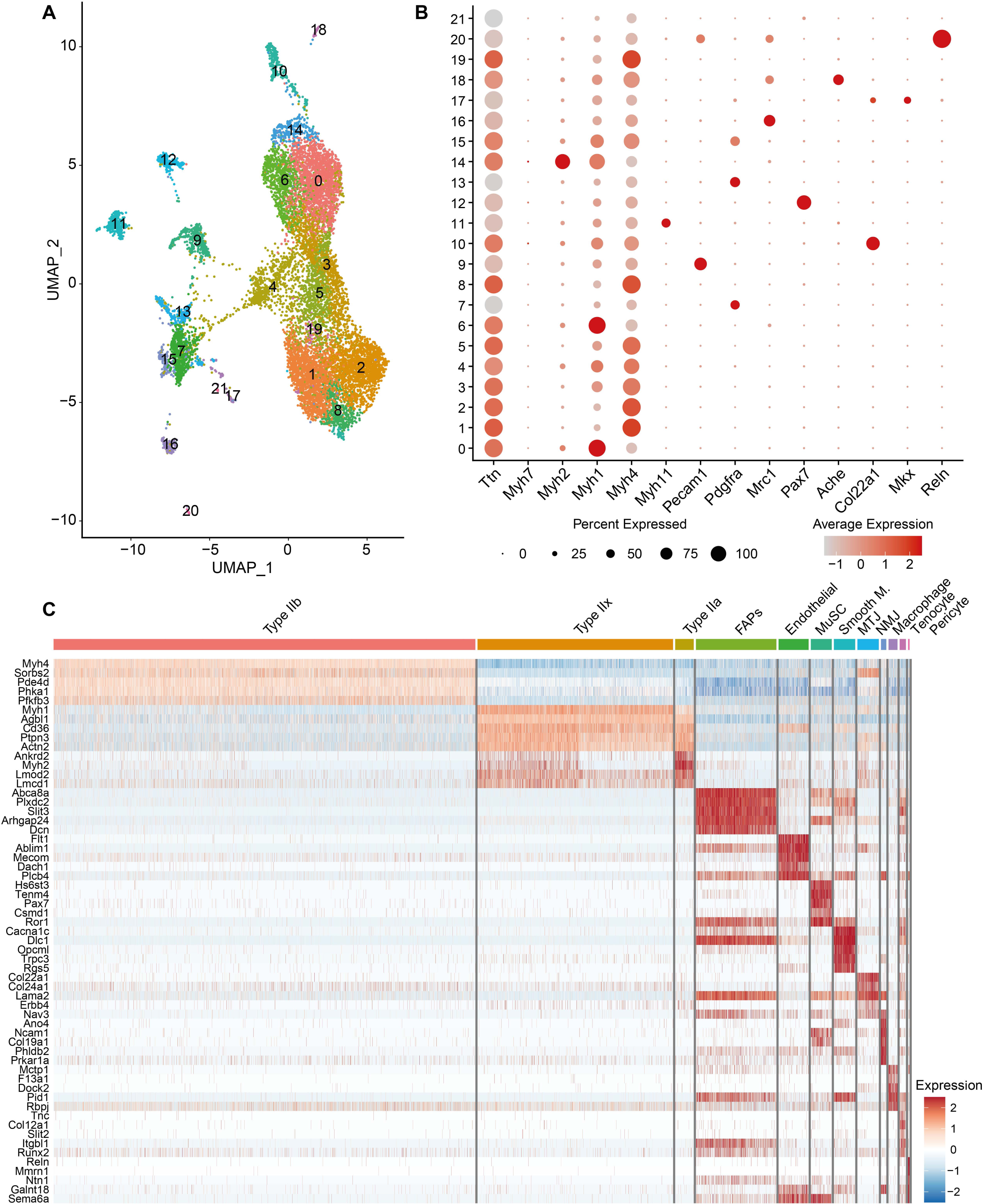
Single-nucleus RNA-seq analysis of atrophying muscles identifies distinct nuclear signatures. (A) UMAP plot of color-coded unsupervised clusters. (B) Dot plot of marker gene expression in unsupervised clusters. The size of the dots represents the percentage of nuclei expressing the marker gene and the red color intensity indicates the expression level. (C) Heatmap of the top 5 signature genes of each nuclear cluster.

**Figure S2.**
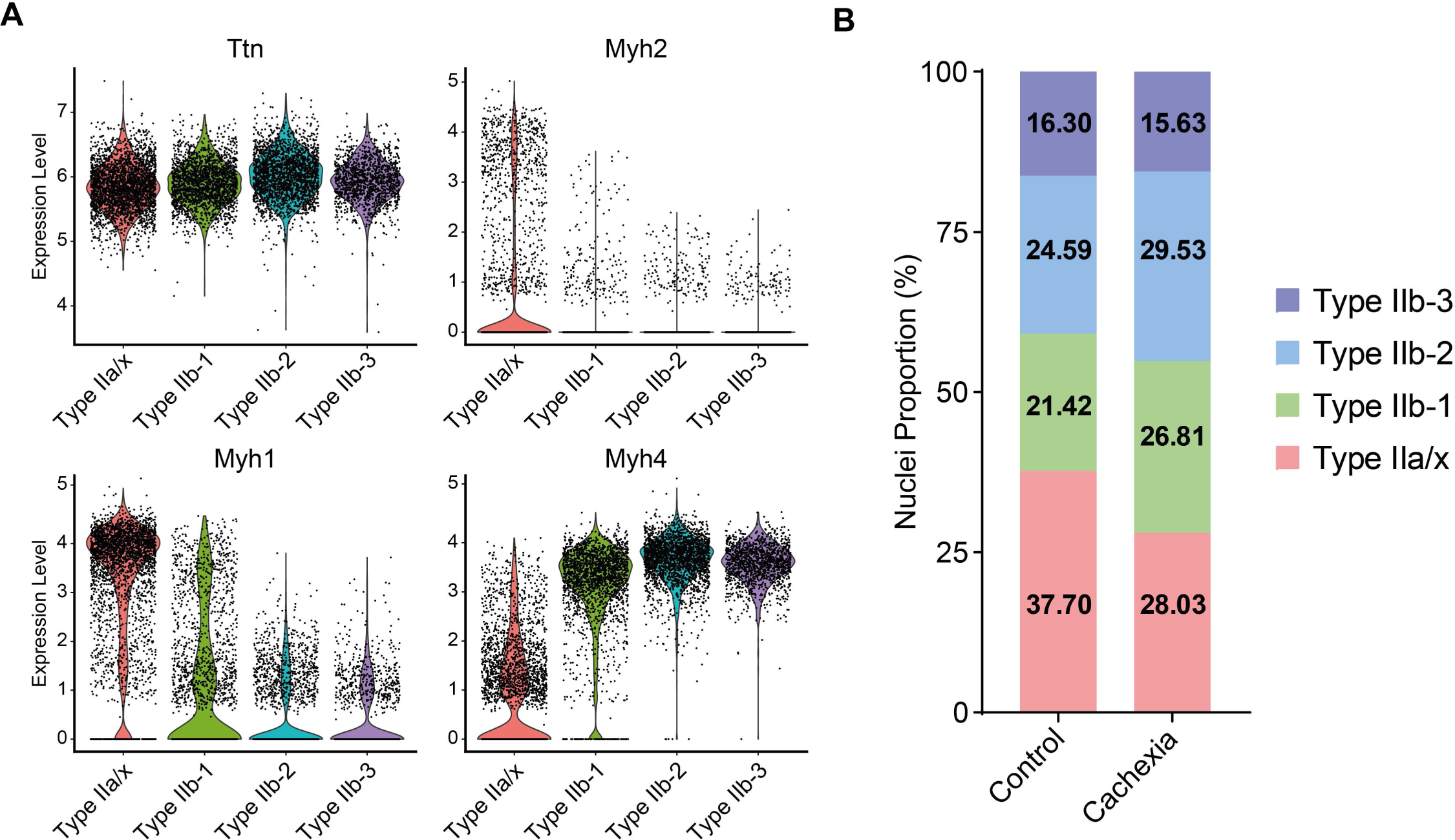
Tumors induce the enrichment of type IIb myonuclear signatures in TA muscle. (A) Violin plot of expression levels of the selected genes in myonuclear subtypes. (B) Proportions of myonuclear clusters in control and cachectic muscles.

**Figure S3.**
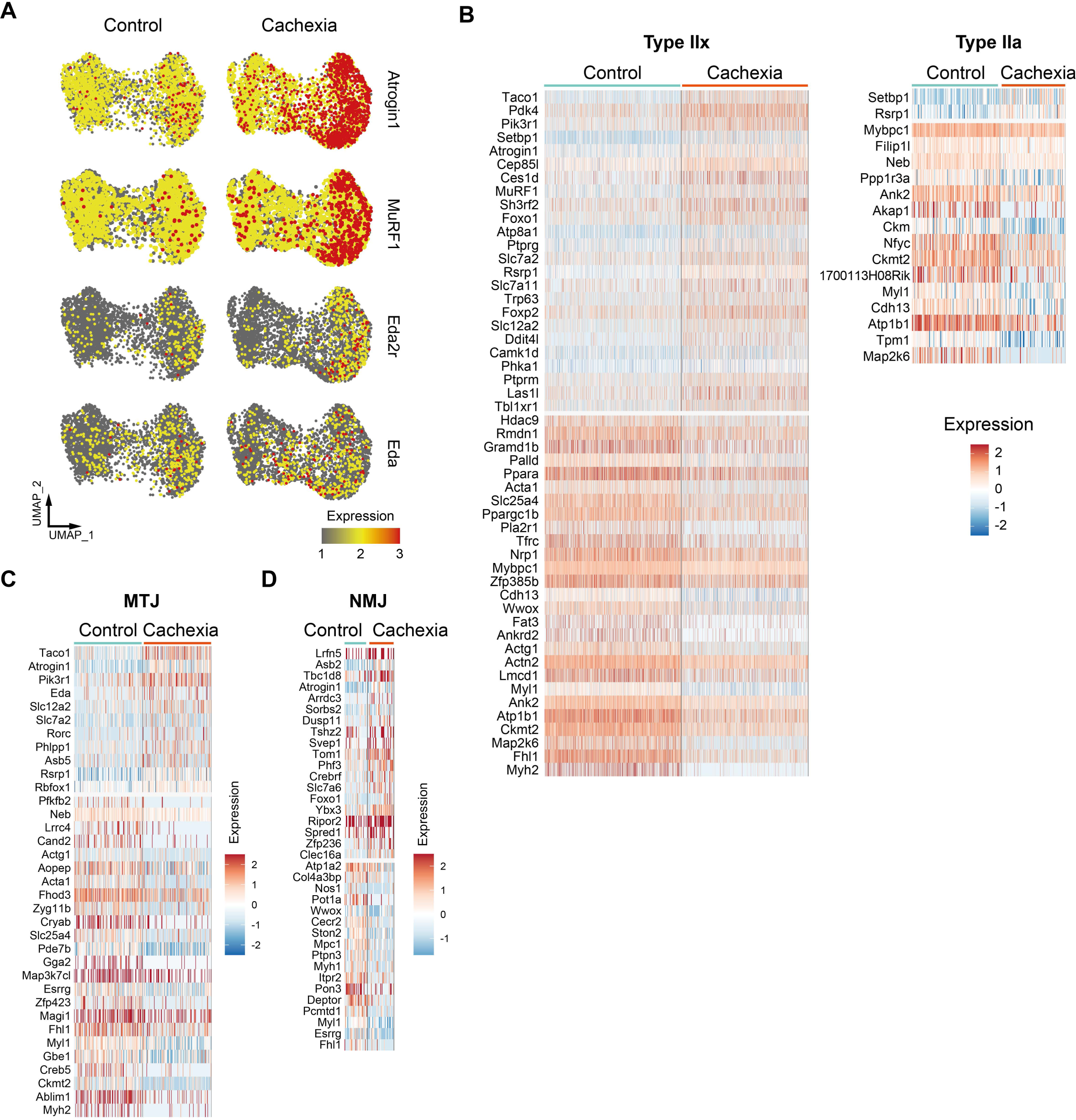
Tumors promote the enrichment of atrophy-related myonuclear signatures in TA muscle. (A) UMAP plots representing expression levels of selected genes in control (left) and cachectic (right) myonuclei. (B) Heatmap of the differentially expressed genes (DEGs) in type IIx (left panel) and type IIa (right panel) myonuclei. (C,D) Heatmap of the DEGs in MTJ (C) and NMJ (D) nuclei.

**Figure S4.**
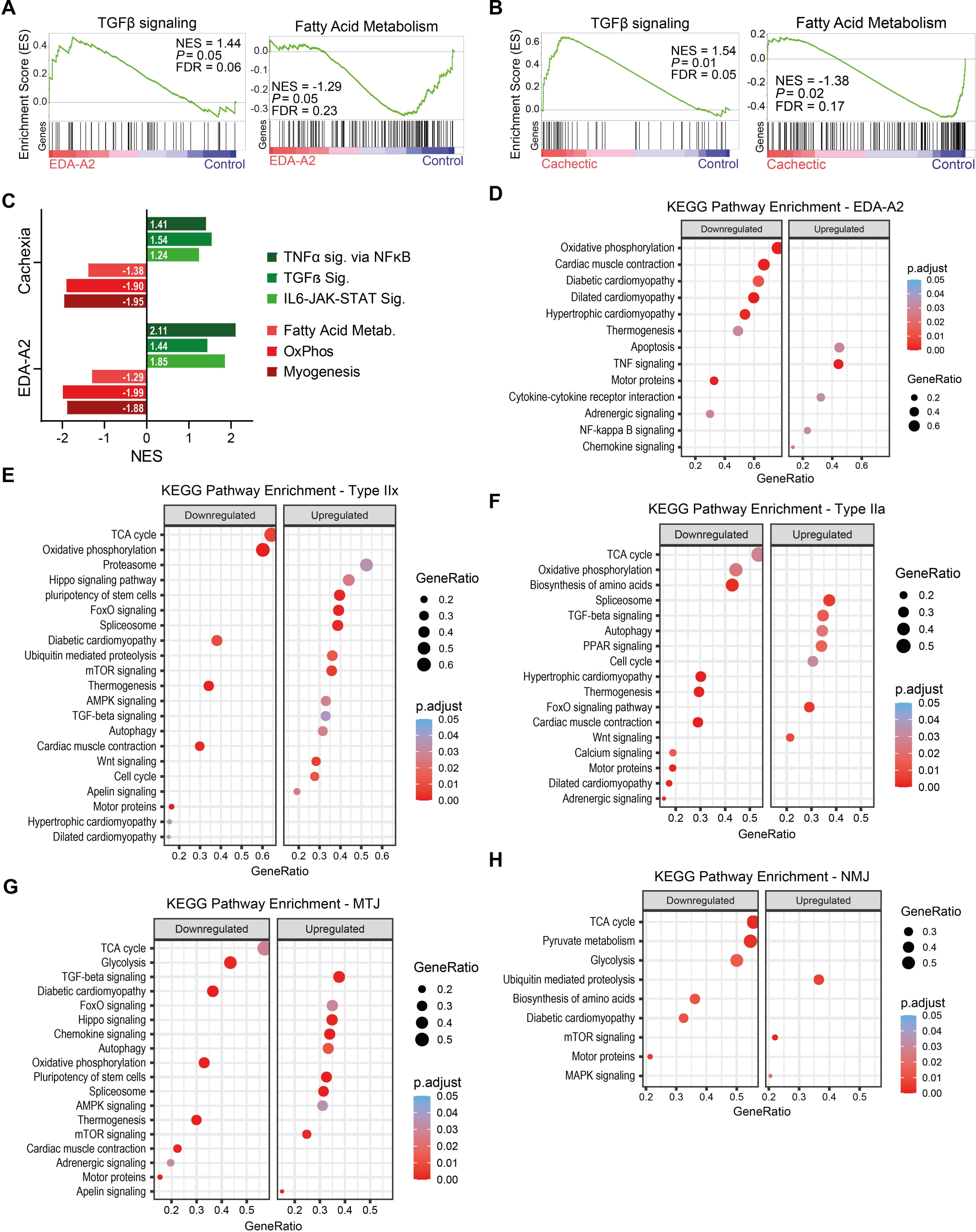
EDA2R activation and tumor inoculation suppress gene sets related to muscle contraction and oxidative metabolism. (A,B) GSEA plots of hallmark gene sets enriched in EDA-A2-treated myotubes (A) and cachectic type IIb myonuclei (B). NES, normalized enrichment score; FDR, false discovery rate. (C) Comparison of NES values of hallmark gene sets enriched in type IIb myonuclei and EDA-A2-treated myotubes. (D-H) KEGG pathways enriched in EDA-A2 treated myotubes (D), cachectic type IIx myonuclei (E), cachectic type IIa myonuclei (F), cachectic MTJ nuclei (G), and cachectic NMJ nuclei (H). The size of the dots represents the enriched gene ratio in each pathway and the red color intensity represents adjusted p values.

**Figure S5.**
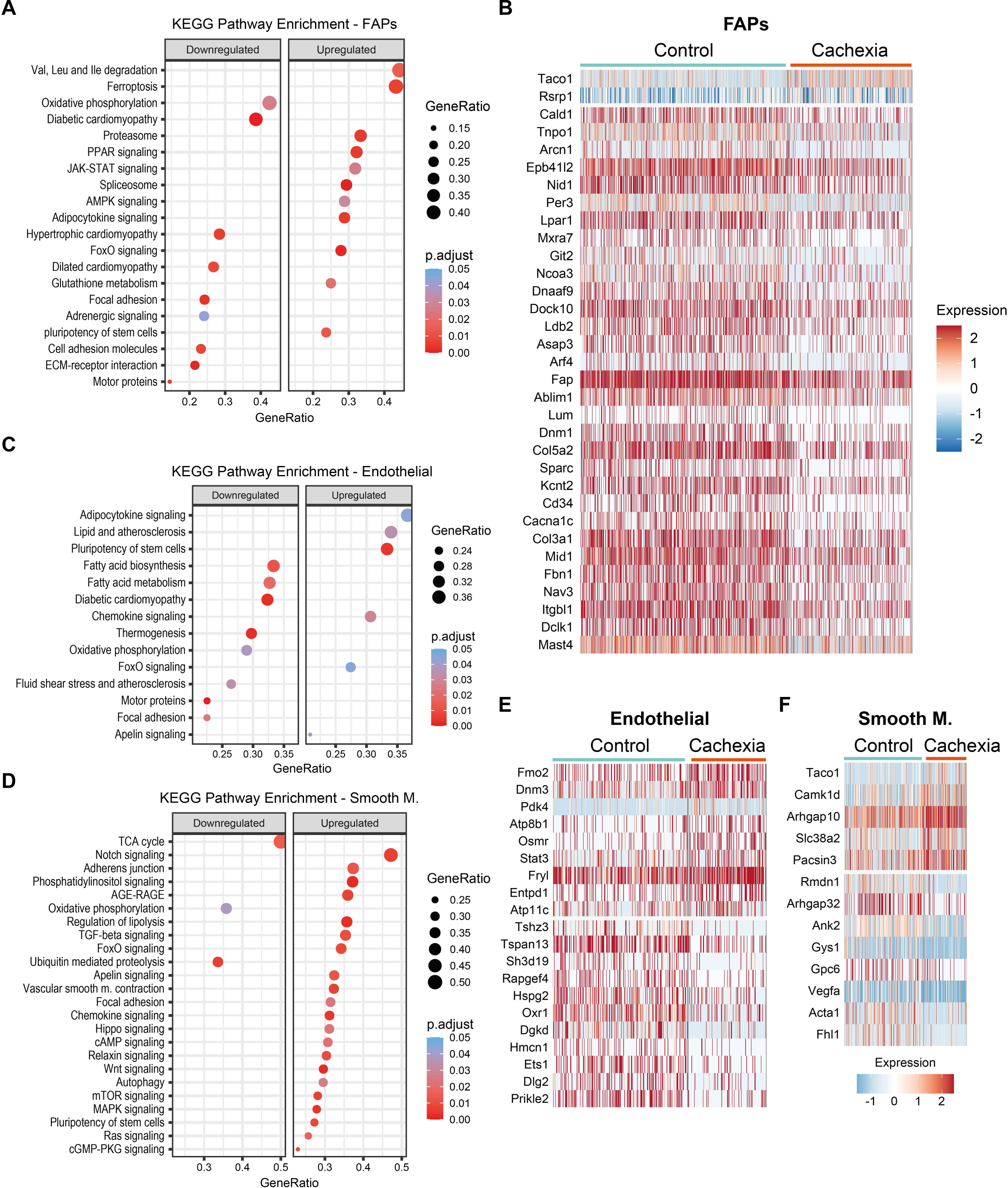
Tumors alter the transcriptomes of mononuclear cells in muscle tissue. (A) KEGG pathways enriched in cachectic FAPs nuclei. (B) Heatmap of the DEGs in FAPs nuclei. (C,D) KEGG pathways enriched in cachectic endothelial nuclei (C) and cachectic smooth muscle nuclei (D). The size of the dots represents the enriched gene ratio in each pathway and the red color intensity represents adjusted p values. (E,F) Heatmap of the DEGs in endothelial nuclei (E) and smooth muscle nuclei (F).

## Notes

### Competing Interest Statement

The authors have declared no competing interest.

